# Co-linear Chaining on Pangenome Graphs

**DOI:** 10.1101/2023.06.21.545871

**Authors:** Jyotshna Rajput, Ghanshyam Chandra, Chirag Jain

## Abstract

Pangenome reference graphs are useful in genomics because they compactly represent the genetic diversity within a species, a capability that linear references lack. However, efficiently aligning sequences to these graphs with complex topology and cycles can be challenging. The seed-chain-extend based alignment algorithms use co-linear chaining as a standard technique to identify a good cluster of exact seed matches that can be combined to form an alignment. Recent works show how the co-linear chaining problem can be efficiently solved for acyclic pangenome graphs by exploiting their small width [Makinen *et al*., TALG’19] and how incorporating gap cost in the scoring function improves alignment accuracy [Chandra and Jain, RECOMB’23]. However, it remains open on how to effectively generalize these techniques for general pangenome graphs which contain cycles. Here we present the first practical formulation and an exact algorithm for co-linear chaining on cyclic pangenome graphs. We rigorously prove the correctness and computational complexity of the proposed algorithm. We evaluate the empirical performance of our algorithm by aligning simulated long reads from the human genome to a cyclic pangenome graph constructed from 95 publicly available haplotype-resolved human genome assemblies. While the existing heuristic-based algorithms are faster, the proposed algorithm provides a significant advantage in terms of accuracy.

**Implementation:** https://github.com/at-cg/PanAligner

## 1 Introduction

Graph-based representation of genome sequences has emerged as a prominent data structure in genomics, offering a powerful means to represent the genetic variation within a species across multiple individuals [11, 17, 26, 49, 51, 53]. A pangenome graph can be represented as a directed graph *G*(*V, E*) such that vertices are labeled by characters (or strings) from the alphabet {A,C,G,T}. The topology of the graph is determined by the count and the type of variants included in the graph. For example, inversions, duplications, or copy number variation are best represented as cycles in a pangenome graph [8, 26, 27, 41, 49]. As a result, the draft pangenome graphs published by the Human Pangenome Reference Consortium [26] and the Chinese Pangenome Consortium [14] are also cyclic. Aligning reads or assembly contigs to a directed labeled graph is a fundamental problem in computational pangenomics [2, 7]. Aligning reads to graphs is also useful for other bioinformatics tasks such as long-read *de novo* assembly [6, 15, 43] and long-read error correction [28, 47].

Formally, the sequence-to-graph alignment problem seeks a walk in the graph that spells a sequence with minimum edit distance from the input sequence. *O*(|*Q*||*E*|) time alignment algorithms for this problem are already known, where *Q* is the query sequence [22, 36]. The known conditional lower bound [3] implies that an exact algorithm significantly faster than *O*(|*Q*||*E*|) is unlikely. This lower bound also holds for de Bruijn graphs [18]. Therefore, fast heuristics are used to process high-throughput sequencing data.

Seed-chain-extend is a common heuristic used by modern alignment tools [21, 23, 46]. This is a three-step process. First, the seeding stage involves computing exact seed matches, such as *k*-mer matches, between a query sequence and a reference. These matches are referred to as *anchors*. The presence of repetitive sequences in genomes often leads to a large number of false positive anchors. Subsequently, the *chaining* stage is employed to link the subsets of anchors in a coherent manner while optimizing specific criteria (Figure 1). This procedure also eliminates the false positive anchors. Finally, the extend stage returns a base-to-base alignment along the selected anchors. Efficient generalization of the three stages to pangenome graphs is an active research topic [7]. Many sequence-to-graph aligners already exist that differ in terms of implementing these stages [5, 9, 24, 30, 42, 49]. This paper addresses the generalization of the chaining stage to cyclic pangenome graphs.

**Figure 1.**
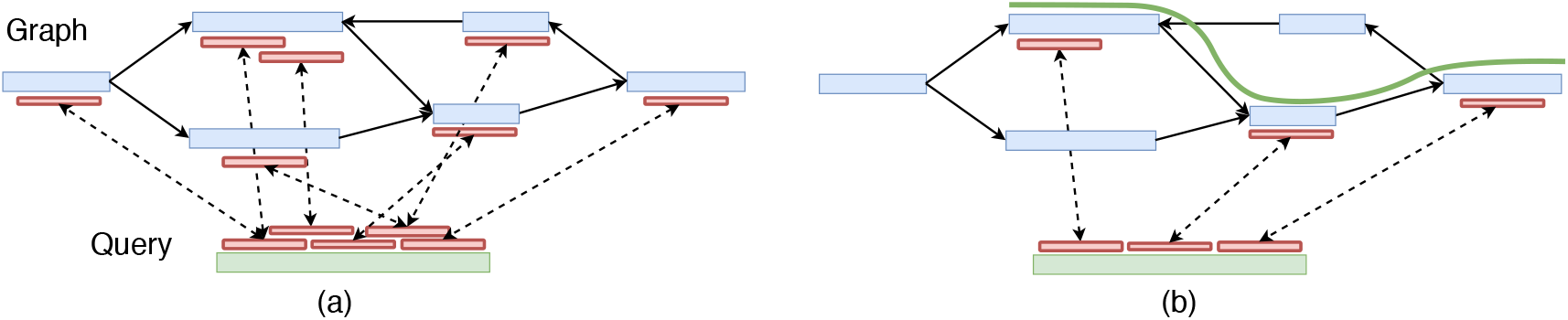
An illustration of co-linear chaining for sequence-to-graph alignment. Assume that the vertices of the graph are labeled with nucleotide sequences. Short exact matches, i.e., anchors, are illustrated using red blocks joined by dotted lines. In (b), the anchors corresponding to the best-scoring chain are retained, and the rest are removed. The retained anchors are combined to produce an alignment of the query sequence to the graph.

**Figure 2.**
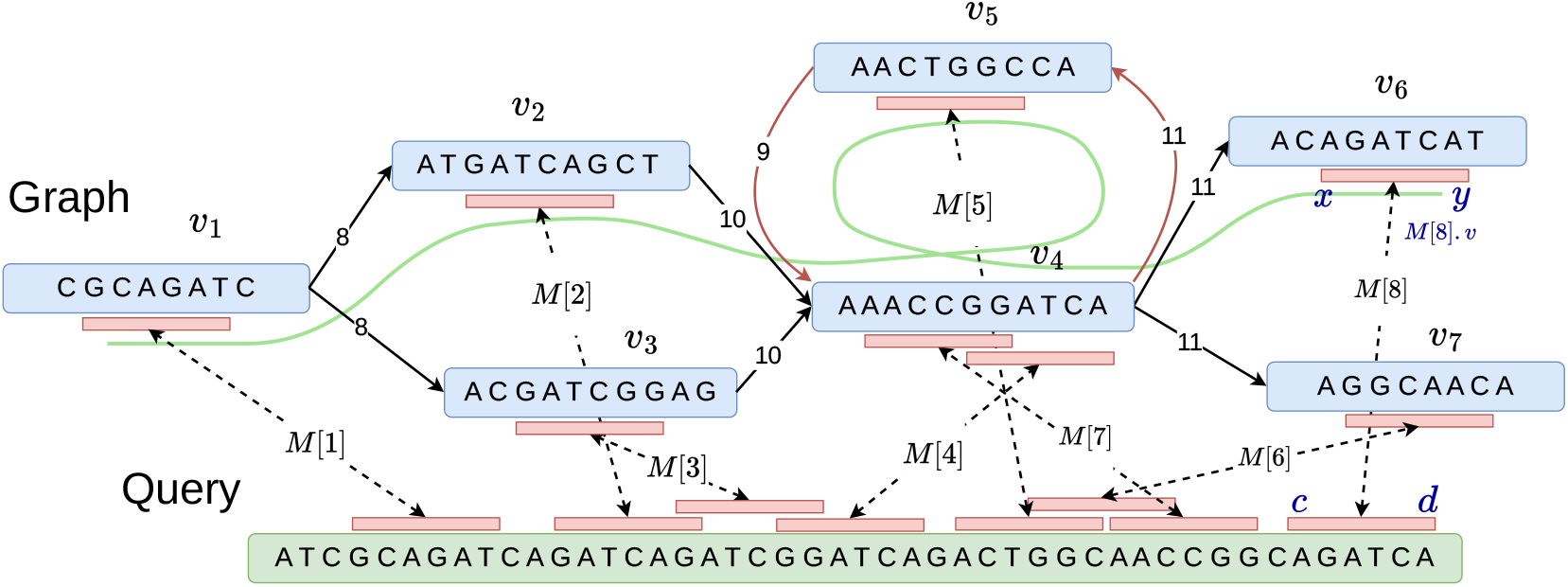
An example illustrating a graph, a query sequence, and multiple anchors as input for co-linear chaining. The sequence of anchors (*M* [1], *M* [2], *M* [4], *M* [5], *M* [7], *M* [8]) forms a valid chain that visits vertex *v*_4_ twice due to a cycle in the graph. The coordinates associated with anchor *M* [8] are also highlighted as an example.

**Figure 3.**
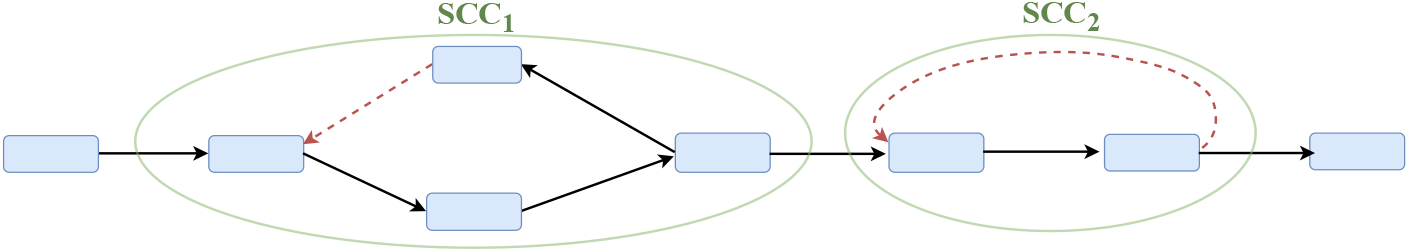
An illustration of the proposed heuristic used to convert a cyclic graph into a DAG. Red-dotted edges represent the removed back edges in each strongly connected component (SCC).

### 1.1 Related Work

Co-linear chaining is a mathematically rigorous method to filter anchors after the seeding stage. It has been well-studied for the sequence-to-sequence alignment case [1, 12, 13, 20, 32, 35, 40]. The input to the chaining problem is a set of *N* weighted anchors. An anchor can be denoted as a pair of intervals in the two sequences corresponding to the exact seed match. A chain is an ordered subset of anchors whose intervals must appear in increasing order in both sequences. The co-linear chaining problem seeks the chain with the highest score, where the score of a chain is calculated by summing the weights of the anchors in the chain and subtracting the penalty for gaps between adjacent anchors. The problem is solvable in *O*(*N* log *N*) time [1].

The first effort to generalize the co-linear chaining problem to graphs was made by Makinen *et al*. [33]. They addressed the co-linear chaining problem on directed acyclic graphs (DAGs). The authors introduced a sparse dynamic programming algorithm whose runtime complexity is parameterized in terms of the *width* of the DAG. The width of a DAG is defined as the minimum number of paths in the DAG such that each vertex is included in at least one path. Parameterizing the complexity in terms of the width is helpful because pangenome graphs typically have small width in practice [5, 30, 33]. An optimized version of their algorithm requires *O*(*KN* log *KN*) time for chaining, where *K* is the width of the DAG [30]. This formulation has been further extended to incorporate gap cost in the scoring function [5], and for solving the longest common subsequence problem between a DAG and a sequence [44]. However, there is limited work on formulating and solving the co-linear chaining problem for general pangenome graphs which might contain cycles. One way to address this was discussed in [30, Appendix section], but the proposed formulation is oblivious to the coordinates of anchors that lie in a strongly connected component of the graph. Their algorithm works by shrinking every strongly connected component into a single vertex and applying the same algorithm developed for DAGs. With this approach, the high-scoring anchor chains in cyclic regions of the graph may result in low-quality alignments.

### 1.2 Contributions

In this paper, we build on top of the algorithmic techniques developed for DAGs [5, 30, 33] and propose novel formulations for cyclic pangenome graphs. Our proposed algorithm exploits the small width of pangenome graphs similar to [33]. Our approach for defining the gap cost between a pair of anchors is inspired by the corresponding function defined on DAGs [5].

We address the following three challenges that arise on cyclic pangenome graphs. First, the dynamic programming-based chaining algorithms developed for DAGs exploit the topological ordering of vertices [5, 30, 33], but such an ordering is not available in cyclic graphs. Second, computing the width and a minimum path cover can be solved in polynomial time for DAGs but is NP-hard for general instances [4]. Third, the walk corresponding to the optimal sequence-to-graph alignment can traverse a vertex multiple times if there are cycles. Accordingly, a chain of anchors should be allowed to loop through vertices. Our proposed problem formulation and the proposed algorithm address the above challenges. Our approach involves computing a path cover 𝒫 of the input graph followed by using iterative algorithms. Let Γ_*c*_, Γ_*l*_, Γ_*d*_ be the parameters that specify the count of iterations used in our algorithms (formally defined later). Our chaining algorithm solves the stated objective in *O*(Γ_*c*_|𝒫|*N* log *N* + |𝒫|*N* log |𝒫|*N*) time after a one-time preprocessing of the graph in *O*((Γ_*l*_ + Γ_*d*_ + log |*V* |)|𝒫||*E*|) time. We will show that parameters |𝒫|, Γ_*c*_, Γ_*l*_, Γ_*d*_ are small in practice to justify the practicality of this approach.

We implemented the proposed chaining algorithm as an open-source software PanAligner. We designed PanAligner as an end-to-end sequence-to-graph aligner using seeding and alignment code from Minigraph [24]. We evaluated the scalability and alignment accuracy of PanAligner by using a cyclic human pangenome graph constructed from 94 high-quality haplotype-resolved assemblies [26] and CHM13 human genome assembly [38]. We achieve the highest long-read mapping accuracy 98.7% using PanAligner when compared to existing methods Minigraph [24] (98.1%) and GraphAligner [42] (97.0%).

## 2 Notations and Problem Formulations

Pangenome graph *G*(*V, E, σ*) is a string labeled graph such that function *σ* : *V →* Σ+ labels each vertex *v* with string *σ*(*v*) over alphabet Σ = *{A, C, G, T }*. Let *Q* be a query sequence over Σ. Let *M* [1..*N*] be an array of anchor tuples (*v*, [*x*..*y*], [*c*..*d*]) with the interpretation that substring *σ*(*v*)[*x*..*y*] from the graph matches substring *Q*[*c*..*d*] in the query sequence. Throughout this paper, all indices start at 1. We will assume that |*E* |≥ |*V* | − 1. Function *weight* assigns a user-specified weight to each anchor. For example, the weight of an anchor could be proportional to the length of the matching substring.

A path cover is a set 𝒫 = *{P*_1_, *P*_2_, …, *P*_|𝒫 |_*}* of paths in graph *G* such that every vertex in *V* is included in at least one of the |𝒫 |paths. We define *paths*(*v*) as *{i* : *P*_*i*_ includes *v}*. If *i ∈ paths*(*v*), then let *index*(*v, i*) specify the position of vertex *v* on path *P*_*i*_. Suppose ℛ^*−*^(*v*) is the set of vertices in *V* that can reach vertex *v* through any walk in graph *G*. We will assume that the set ℛ^*−*^(*v*) always includes the vertex *v*. The value *last*2*reach*(*v, i*) for *v ∈ V, i ∈* [1, |𝒫|] represents the last vertex on path *P*_*i*_ that belongs to set ℛ^*−*^(*v*). Note that *last*2*reach*(*v, i*) does not exist if there is no vertex on path *P*_*i*_ that belongs to ℛ^*−*^(*v*). Let *N* ^+^(*v*) and *N* ^*−*^(*v*) be the set of outgoing and incoming neighbor vertices of vertex *v*, respectively.

We need to calculate character distances between pairs of anchors in the graph while solving the co-linear chaining problem. Assume that edge (*v, u*) *∈ E* has length |*σ*(*v*)|*>* 0. Let *D*(*v*_1_, *v*_2_) denote the length of the shortest path from vertex *v*_1_ to *v*_2_ in *G*. We set *D*(*v*_1_, *v*_2_) = *∞* if there is no path from *v*_1_ to *v*_2_, whereas *D*(*v*_1_, *v*_2_) = 0 if *v*_1_ = *v*_2_. We use *D*^*°*^(*v*) to specify the length of the shortest proper cycle containing *v. D*^*°*^(*v*) = *∞* if *v* is not part of any proper cycle. If *P*_*i*_ includes *v*, let *dist*2*begin*(*v, i*) denote the length of the sub-path of path *P*_*i*_ from the start of *P*_*i*_ to *v*.

Our algorithm will use a balanced binary search tree data structure for executing range queries efficiently. It has the following properties.

### ▸ Lemma 1 (ref. [31]).

*Let n be the number of leaves in a tree, each storing a* (*key, value*) *pair. The following operations can be supported in O*(log *n*) *time:*

- *update*(*k, val*): *For the leaf w with key* = *k, value*(*w*) *←−* max(*value*(*w*), *val*).
- *RMQ*(*l, r*): *Return* max*{value*(*w*) |*l < key*(*w*) *< r} such that w is a leaf. This is the range maximum query*.

*Given n* (*key, value*) *pairs, the tree can be constructed in O*(*n* log *n*) *time and O*(*n*) *space*.

Next, we define a precedence relation between a pair of anchors, which is a partial order among the input anchors [30].

### ▸ Definition 2 (Precedence).

*Given two anchors M* [*i*] *and M* [*j*], *we define M* [*i*] *precedes* (*≺*) *M* [*j*] *as follows. If M* [*i*].*v ?*= *M* [*j*].*v, then M* [*i*] *≺ M* [*j*] *if and only if M* [*i*].*d < M* [*j*].*c and M* [*i*].*v reaches M* [*j*].*v. If M* [*i*].*v* = *M* [*j*].*v, then M* [*i*] *≺ M* [*j*] *if and only if M* [*i*].*d < M* [*j*].*c, and M* [*i*].*y < M* [*j*].*x or the graph has a proper cycle containing M* [*i*].*v*.

### ▸ Definition 3 (Chain).

*Given the set of anchors {M* [1], *M* [2], …, *M* [*N*]*}, a chain is an ordered subset of anchors S* = *s*_1_*s*_2_ *⋯s*_*q*_ *of M, such that s*_*j*_ *precedes s*_*j*+1_ *for all* 1 *≤ j < q*.

Our co-linear chaining problem formulation seeks a chain *S* = *s*_1_*s*_2_ *…s*_*q*_ that maximizes the chain score defined as 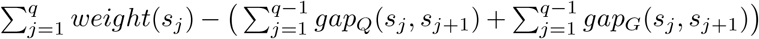. Functions *gap*_*Q*_ and *gap*_*G*_ specify the gap cost incurred in the query sequence and the graph, respectively. Although we specifically focus on problem formulations where the gap cost is calculated as the sum of *gap*_*G*_ and *gap*_*Q*_, our approach can be extended to other gap definitions such as |*gap*_*G*_ *− gap*_*Q*_|, min(*gap*_*G*_, *gap*_*Q*_), or max(*gap*_*G*_, *gap*_*Q*_), similar to [5]. We define *gap*_*Q*_(*s*_*j*_, *s*_*j*+1_) as *s*_*j*+1_.*c − s*_*j*_.*d −* 1, which can be interpreted as the count of characters in the query sequence between the endpoints of the two anchors. Next, we will define two versions of the co-linear chaining problem that differ in their definition of *gap*_*G*_. In both versions, *gap*_*G*_(*s*_*j*_, *s*_*j*+1_) is calculated by looking at the count of characters spelled along a walk in the graph from *s*_*j*_ to *s*_*j*+1_. In the first version of the problem formulation, we use the shortest path from vertex *s*_*j*_.*v* to *s*_*j*+1_.*v* to calculate *gap*_*G*_(*s*_*j*_, *s*_*j*+1_).

### ▸ Problem 4.

*Given a query sequence Q, graph G*(*V, E, σ*) *and anchors M* [1..*N*], *determine the optimal chaining score by using the following definition of gap*_*G*_:

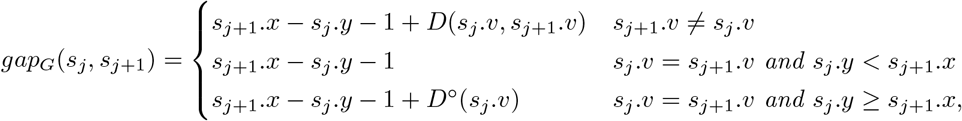

*where* (*s*_*j*_, *s*_*j*+1_) *is a pair of anchors from M such that s*_*j*_ *precedes s*_*j*+1_.

### ▸ Lemma 5.

*Problem 4 can be solved in* Θ(|*V* ||*E*|+ |*V* |^2^ log |*V* |+ *N* ^2^) *time*.

Proof. Compute the shortest distance *D*(*v*_*i*_, *v*_*j*_) between all pairs of vertices *v*_*i*_, *v*_*j*_ *∈ V* in *O*(|*V* ||*E*|+ |*V* |^2^ log |*V* |) time by using Dijkstra’s algorithm from every vertex. Next, compute *D*^*°*^(*v*) as min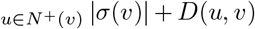 in Θ(|*E*|) time for all *v ∈ V*. These computations need to be done only once for a graph. To solve the chaining problem for a given query sequence, sort the input anchor array *M* [1..*N*] in non-decreasing order by the component *M* [·].*c*. Let *C*[1..*N*] be a one-dimensional table in which *C*[*j*] will be the optimal score of a chain ending at anchor *M* [*j*]. Initialize *C*[*j*] as *weight*(*M* [*j*]) for all *j ∈* [1, *N*]. Subsequently, compute *C* in the left-to-right order by using the recursion *C*[*j*] = max_*M*[*i*]*≺M*[*j*]_*{C*[*j*], *weight*(*M* [*j*]) *− gap*_*Q*_(*M* [*i*], *M* [*j*]) *− gap*_*G*_(*M* [*i*], *M* [*j*])*}*. Computing *C*[*j*] takes Θ(*N*) time because precedence condition can be checked in constant time. Report max_*j*_ *C*[*j*] as the optimal chaining score. ◂

The above algorithm is unlikely to scale to large whole-genome sequencing datasets because it requires Θ(*N* ^2^) time for the dynamic programming recursion. Motivated by [5], we will define an alternative definition of *gap*_*G*_. We will approximate the distance between a pair of vertices by using a path cover of the graph. We will later propose an efficient algorithm for the revised problem formulation.

Suppose 𝒫 = *{P*_1_, *P*_2_, …, *P*_|𝒫|_*}* is a path cover of graph *G*. Consider a pair of vertices *v*_1_, *v*_2_ *∈ V* such that *v*_1_ reaches *v*_2_. For each path *i ∈ paths*(*v*_1_), consider the walk starting from *v*_1_ along the edges of path *P*_*i*_ till vertex *α*_*i*_, where vertex *α*_*i*_ = *v*_2_ if *v*_2_ also lies on path *P*_*i*_ anywhere after *v*_1_, i.e., *index*(*v*_2_, *i*) ≥ *index*(*v*_1_, *i*), and *α*_*i*_ = *last*2*reach*(*v*_2_, *i*) otherwise.

If *α*_*i*_≠ *v*_2_, the rest of the walk till *v*_2_ is completed by using the shortest path from vertex *α*_*i*_ to *v*_2_. Denote *D*_𝒫_ (*v*_1_, *v*_2_) as the length of the shortest walk among such |*paths*(*v*_1_)|possible walks from *v*_1_ to *v*_2_. Formally, we can write *D*_𝒫_ (*v*_1_, *v*_2_) as

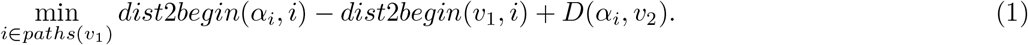

*D*_𝒫_ (*v*_1_, *v*_2_) is well defined if *v*_2_ is reachable from *v*_1_. We set *D*_𝒫_ (*v*_1_, *v*_2_) = *∞* if *v*_2_ is not reachable from *v*_1_. Finally, if vertex *v* is part of a proper cycle in *G*, we define 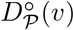 as the length of a specific walk that starts and ends at *v*, i.e., 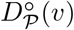 as min 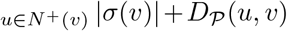 for all 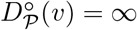 if *v* is not part of any proper cycle.

### ▸ Problem 6.

*Given a query sequence Q, graph G*(*V, E, σ*) *and anchors M* [1..*N*], *determine a path cover P of the graph, and the optimal chaining score by using the following definition of gap*_*G*_:

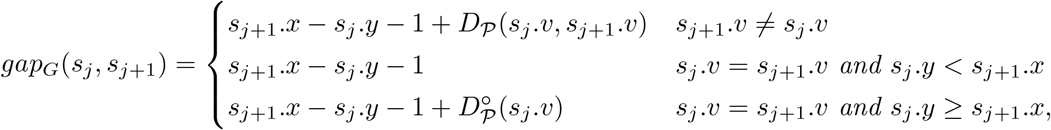

*where* (*s*_*j*_, *s*_*j*+1_) *is a pair of anchors from M such that s*_*j*_ *precedes s*_*j*+1_.

## 3 Proposed Algorithms

A single experiment typically requires aligning millions of reads to a graph. Therefore, we will do a one-time preprocessing of the graph that will help reduce the runtime of our chaining algorithm for solving Problem 6.

### 3.1 Algorithms for Preprocessing the Graph

We compute the following quantities during the preprocessing stage:

▪ A path cover 𝒫 of *G*(*V, E, σ*). We require the path cover to be small (in the number of paths). However, determining the minimum path cover in a graph with cycles is an *NP* -hard problem. We will discuss an efficient heuristic for determining a small path cover.
▪ A bijective function *rank* : *V →* [1, |*V* |] that specifies a linear ordering of vertices. The ordering should satisfy the following property: If vertex *v*_2_ occurs anywhere after *v*_1_ in a path in 𝒫, then *rank*(*v*_2_) *> rank*(*v*_1_) for all *v*_1_, *v*_2_ *∈ V*. Such an ordering may not exist for an arbitrary path cover but it will exist for the path cover chosen by us.
▪ *last*2*reach*(*v, i*), *D*(*last*2*reach*(*v, i*), *v*), *dist*2*begin*(*v, i*) and 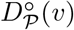 for all *v ∈ V* and *i ∈* [1, |𝒫|]. These values will be frequently used by our chaining algorithm to compute gap costs.

We propose the following heuristic for computing a small path cover of graph *G*(*V, E, σ*). We derive a DAG *G*^*′*^(*V, E*^*′*^, *σ*) from *G* by removing a small number of edges. Next, we determine the minimum path cover 𝒫 of *G*^*′*^ in *O*(|𝒫||*E*|log |*V* |) time by using a known algorithm [33]. Our intuition is that removing as few edges as possible will provide a close to optimal path cover of *G*. One way to compute *G*^*′*^ is to use standard heuristic-based solvers for minimum feedback arc set (FAS) problem, e.g., [10], but we empirically observed that this approach could sometimes disconnect a weak component of a graph, leading to a large path cover. Therefore, instead of using FAS heuristics, we use a simple idea where we identify all strongly connected components [50] and perform a depth-first search within each strong component to remove back edges. This approach provides a DAG that has the same number of weak components as *G* while removing a small number of edges in practice, thus resulting in a small path cover. Next, we compute a function *rank* for all vertices *∈ V* by topological sorting of vertices in DAG *G*^*′*^.

If there is no cycle in *G*, then *last*2*reach*(*v, i*) and *D*(*last*2*reach*(*v, i*), *v*) can be computed in *O*(|𝒫||*E*|) time by using dynamic programming algorithms that process vertices in topological order [5, 33]. We extend these ideas to cyclic graphs by designing iterative algorithms. We will formally prove that as the iterations proceed, the output gets closer to the desired solution. Our approach to computing *last*2*reach*(*v, i*) is outlined in Algorithm 1. If *last*2*reach*(*v, i*) exists, the algorithm determines it in terms of its *rank*. We maintain an array *L*2*R* to save intermediate results. *L*2*R*(*v, i*) is initialised to *rank*(*v*) if *v* lies on path *P*_*i*_. In each iteration, we revise *L*2*R*(*v, i*) by probing *L*2*R*(*u, i*) for all *u ∈ N* ^*−*^(*v*). In Lemma 7, we prove the correctness of this algorithm by arguing that all |𝒫||*V* |values in array *L*2*R* converge to their optimal values through label propagation in *≤* |*V* |iterations. Let Γ_*l*_ denote the count of iterations used by the algorithm. *L*2*R*(*v, i*) remains 0 if *last*2*reach*(*v, i*) does not exist.

#### Algorithm 1

*O*(Γ_*l*_||𝒫|*E*|) time algorithm to compute *last*2*reach*(*v, i*) for all *v ∈ V* and *i ∈* [1, |𝒫|]

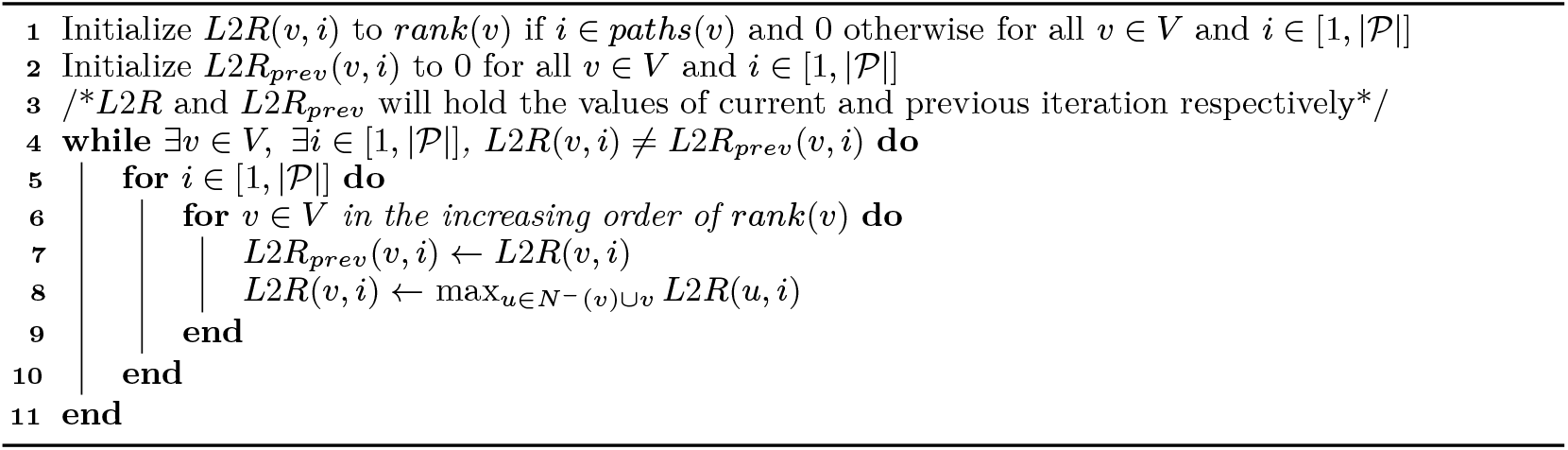

#### ▸ Lemma 7.

*In Algorithm 1, L*2*R*(*v, i*) *converges to the rank of last*2*reach*(*v, i*) *in at most* |*V* |*iterations for all v ∈ V and i ∈* [1, |𝒫|].

Proof. A vertex *v*_2_ *∈ V* is said to be reachable within *k* hops from vertex *v*_1_ *∈ V* if there exists a path with *≤ k* edges from *v*_1_ to *v*_2_. We will prove by induction that Algorithm 1 satisfies the following invariant: After *j* iterations, *L*2*R*(*v, i*) has converged to *rank*(*last*2*reach*(*v, i*)) if *last*2*reach*(*v, i*) exists and vertex *v* is reachable within *j* hops from *last*2*reach*(*v, i*) in *G*. This argument will prove the lemma because vertex *v*_2_ *∈ V* must be reachable within |*V* | − 1 hops from *v*_1_ *∈ V* if *v*_2_ is reachable from *v*_1_. Base case (*j* = 0) holds due to initialisation of *L*2*R*(*v, i*) in Line 1. If *v* lies 0-hop from *last*2*reach*(*v, i*), i.e., *v* = *last*2*reach*(*v, i*), then *v* must lie on path *P*_*i*_ and *rank*(*last*2*reach*(*v, i*)) = *rank*(*v*). Next, assume that the invariant is true for *j* = *n*. Now consider the situation after *n* + 1 iterations. Suppose *v ∈ V* is reachable within *n* + 1 hops from *last*2*reach*(*v, i*). Then, at least one neighbour *u ∈ N* ^*−*^(*v*) of vertex *v* exists which is reachable within *n* hops from *last*2*reach*(*v, i*) and *last*2*reach*(*u, i*) = *last*2*reach*(*v, i*). Based on our assumption, *L*2*R*(*u, i*) must have already converged to *rank*(*last*2*reach*(*u, i*)) before (*n* +1)^*th*^ iteration. Therefore, Line 8 in Algorithm 1 ensures that *L*2*R*(*v, i*) *← rank*(*last*2*reach*(*v, i*)) after (*n* + 1)^*th*^ iteration. ◂

It is possible to design an adversarial example where the algorithm uses Ω(|*V* |) iterations. However, in practice, we expect the algorithm to converge quickly. Each iteration of Algorithm 1 requires *O*(|𝒫||*E*|) time. Therefore, the total worst-case time of Algorithm 1 is bounded by *O*(Γ_*l*_|𝒫||*E*|). A similar approach is applicable to compute *D*(*last*2*reach*(*v, i*), *v*) for all *v ∈ V* and *i ∈* [1, |𝒫|] (Algorithm 2). We use Γ_*d*_ to denote the count of iterations needed in Algorithm 2. Similar to parameter Γ_*l*_ in Algorithm 1, Γ_*d*_ is also upper bounded by |*V* |. We will later show empirically that Γ_*l*_ *≪* |*V* |and Γ_*d*_ *≪* |*V* |in practice.

#### Algorithm 2

*O*(Γ_*d*_|𝒫||*E*|) time algorithm to compute *D*(*last*2*reach*(*v, i*), *v*) for all *v ∈ V* and *i ∈* [1, |𝒫|]

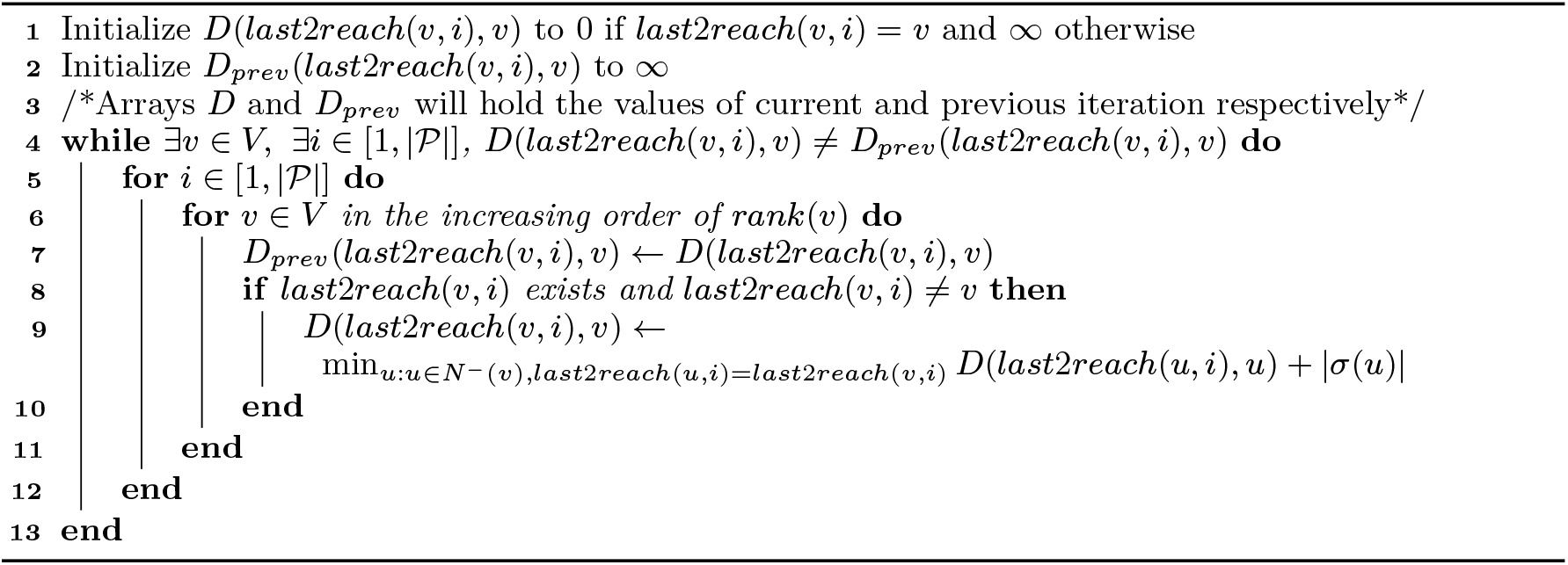

Array *dist*2*begin* is trivially precomputed in *O*(|𝒫||*V* |) time. 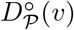 is computed as min 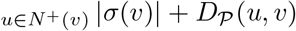 based on its definition. *D*𝒫 (*u, v*) can be calculated by using Equation 1 for any *u, v ∈ V* in *O*(|𝒫|) time. Accordingly, computation of 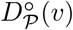 for all *v ∈ V* is done in *O*(|𝒫||*E*|) time. The following lemma summarises the worst-case time complexity of all the preprocessing steps.

#### ▸ Lemma 8.

*Preprocessing of graph G*(*V, E, σ*) *requires O*((Γ_*l*_ + Γ_*d*_ + log |*V* |)|*P*||*E*|) *time*.

### 3.2 Co-linear Chaining Algorithm

We propose an iterative chaining algorithm to address Problem 6. The proposed algorithm builds on top of the known algorithms for DAGs [5, 33]. Similar to [33], we maintain one search tree 𝒯_*i*_ for each path *P*_*i*_ *∈* 𝒫. Given anchors *M* [1..*N*], our algorithm will return array *C*[1..*N*] such that *C*[*j*] corresponds to the optimal score of a chain that ends at anchor *M* [*j*]. If there are no cycles in *G*, then one iteration of Algorithm 3 suffices to compute the optimal chaining scores. For a DAG, a single iteration of Algorithm 3 works equivalently to the known algorithm for DAGs in [5]. In this case, Algorithm 3 would essentially visit the vertices of graph *G* in topological order while ensuring that *C*[*j*] is optimally solved after *M* [*j*].*v* is visited. To solve the chaining problem on cyclic graphs, we design an iterative solution where chaining scores *C*[1..*N*] get closer to optimal values in each iteration. We will use Γ_*c*_ to specify the total count of iterations.

#### Algorithm 3

*O*(Γ_*c*_*N* |𝒫|log *N* + *N* |𝒫|log *N* |𝒫|) time chaining algorithm

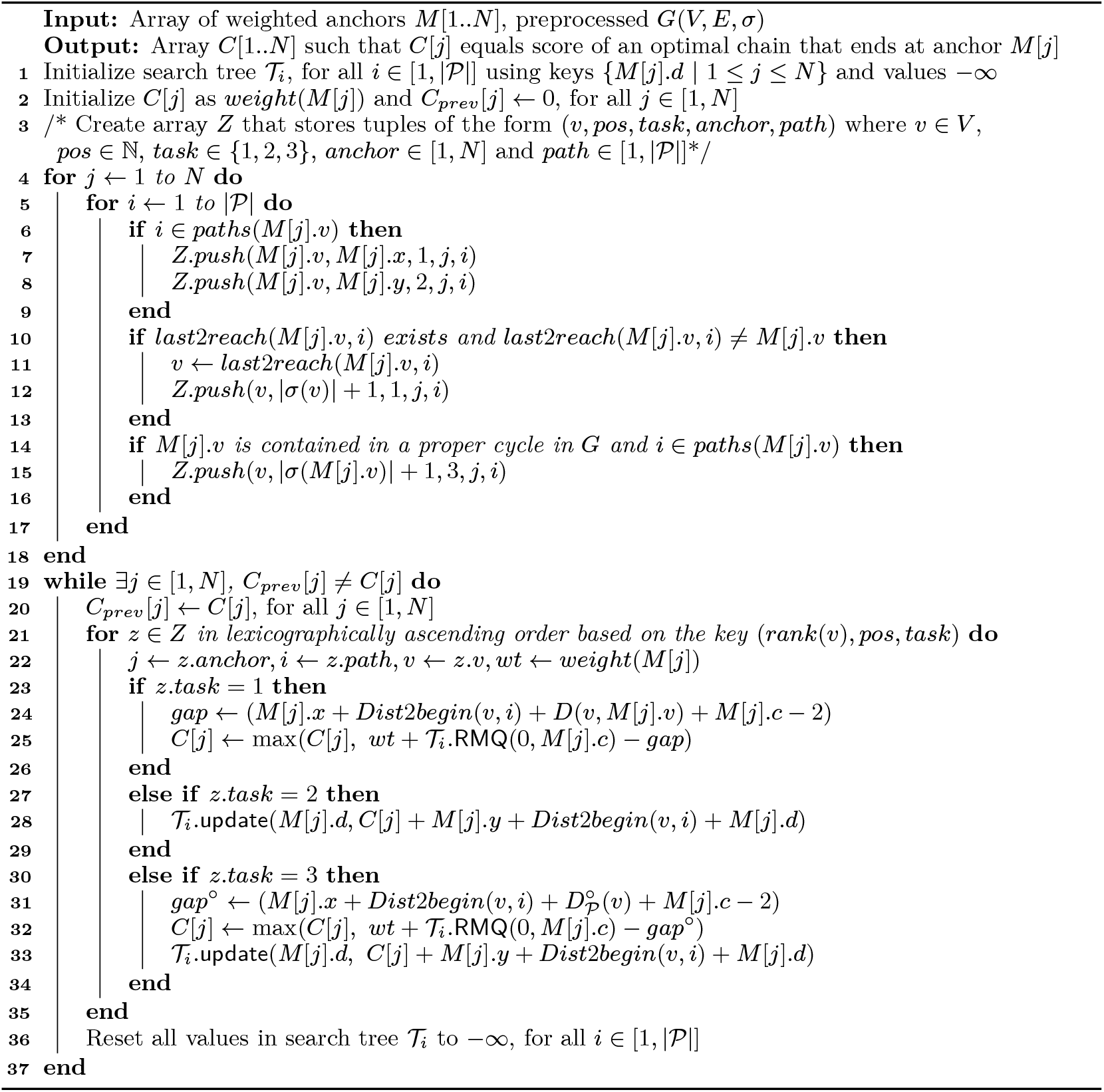

An overview of Algorithm 3 is as follows. At the beginning of each iteration, all search trees 𝒯_*i*_ s are filled with keys *{M* [*j*].*d* |1 *≤ j ≤ N }* and values *−∞*. The values will be used to specify the priorities of anchors based on their scores *C* [ ] and coordinates. Each iteration of our algorithm processes *v ∈ V* in the increasing order of *rank*(*v*). While processing *v*, Algorithm 3 performs three types of tasks:

1. The first type of task is to revise chaining scores *{C*[*j*] : *M* [*j*].*v* = *v}* corresponding to the anchors that lie on vertex *v*. We also revise scores corresponding to those anchors that are located on vertex *u* ≠ *v* such that *v* is the last vertex on a path *∈* 𝒫 to reach *u*. This is achieved by querying search trees 𝒯_*i*_ for all *i ∈ paths*(*v*). In all these tasks, we use *D*𝒫 (*v*_1_, *v*_2_) to calculate distance from vertex *v*_1_ *∈ V* to vertex *v*_2_ *∈ V*.

2. Suppose score *C*[*j*] is revised by using the first category tasks. The second type of task is to update the value of key *M* [*j*].*d* in search trees 𝒯_*i*_ for all *i ∈ paths*(*v*). The value gets updated if the new value is greater than the previously stored value (Lemma 1).

3. The third type of task is to again update scores *{C*[*j*] : *M* [*j*].*v* = *v}* and search trees if *v* is part of a proper cycle in *G*. Here we use *D*^*°*^ (*v*) to calculate the distance of vertex *v* to itself while determining gap costs.

Lines 4-18 in Algorithm 3 build array *Z* that contains up to 4*N* |𝒫|tuples corresponding to all the above type of tasks. Array *Z* is sorted in *O*(*N* |𝒫|log *N* |*P*|) time to ensure that all tasks are executed in the proper order (Line 21). Next, we start the iterative procedure. Lines 19-33 form a single iteration of the algorithm. These tasks lead to updates on score array *C* and the search trees. The arithmetic operations in Lines 24, 25, 31, 32 enable calculation of gap cost based on our definitions of *gap*_*G*_ and *gap*_*Q*_ in Section 2. Each iteration requires *O*(*N* |𝒫|log *N*) time because each task corresponds to either update or RMQ operation on a search tree of size *≤ N*. In Lemma 9, we prove that array *C*[1..*N*] converges to optimality in at most *N* iterations. We will also prove that Ω(*N*) iterations are required for convergence in the worst case.

#### ▸ Lemma 9.

*In Algorithm 3, co-linear chaining scores C*[1..*N*] *converge to optimality in ≤ N iterations*.

Proof. *C*[*j*] always specifies the score of a chain of size ≥ 1 that ends at anchor *M* [*j*] throughout the execution of the algorithm. Let *f*_*i*_(*j*) denote the optimal chaining score ending at anchor *M* [*j*] over all chains of size *≤ i*. We will prove by induction that before *i*^*th*^ iteration begins, *C*[*j*] ≥ *f*_*i*_(*j*) for all *j ∈* [1, *N*]. It suffices to prove this statement because the size of a chain must be *≤ N*. Base case (*i* = 1) holds due to the initialization step in Line 2. Next, assume that before *x*^*th*^ iteration begins, *C*[*j*] ≥ *f*_*x*_(*j*) holds for all *j ∈* [1, *N*]. We will prove that the invariant holds for iteration *x* + 1.

Let *C*_*x*_[*j*] and *C*_*x*+1_[*j*] denote the intermediate values of *C*[*j*] at the start of *x*^*th*^ and (*x*+1)^*th*^ iteration, respectively. From Lines 25 and 32, we know *C*_*x*_[*j*] *≤ C*_*x*+1_[*j*]. If *f*_*x*+1_(*j*) = *f*_*x*_(*j*), then *C*_*x*+1_[*j*] ≥ *C*_*x*_[*j*] ≥ *f*_*x*_(*j*) = *f*_*x*+1_(*j*). Next consider the other case when *f*_*x*+1_(*j*) *> f*_*x*_(*j*). Suppose the optimal chain corresponding to *f*_*x*+1_(*j*) is *M* [*β*_1_], *M* [*β*_2_], …, *M* [*β*_*x*_], *M* [*j*] where *β*_*i*_ *∈* [1, *N*] for all *i ∈* [1, *x*]. Accordingly, *f*_*x*+1_(*j*) = *weight*(*M* [*j*]) + *f*_*x*_(*β*_*x*_) *− gap*_*Q*_(*M* [*β*_*x*_], *M* [*j*]) *− gap*_*G*_(*M* [*β*_*x*_], *M* [*j*]). Based on our induction hypothesis, *C*[*β*_*x*_] ≥ *f*_*x*_(*β*_*x*_) at the start of the *x*^*th*^ iteration. Each iteration of Algorithm 3 processes *v ∈ V* by increasing the order of *rank*(*v*). To prove that *C*_*x*+1_[*j*] ≥ *f*_*x*+1_(*j*), we have the following four cases to consider:

▪ Case 1: *rank*(*M* [*β*_*x*_].*v*) *< rank*(*M* [*j*].*v*). The algorithm processes vertex *M* [*β*_*x*_].*v* before vertex *M* [*j*].*v*. When *M* [*β*_*x*_].*v* is processed during the *x*^*th*^ iteration, the value of key *M* [*β*_*x*_].*d* gets updated in search trees (Line 28). *C*[*j*] gets updated later. At the end of the *x*^*th*^ iteration, *C*[*j*] ≥ *weight*(*M* [*j*]) + *f*_*x*_(*β*_*x*_) *− gap*_*Q*_(*M* [*β*_*x*_], *M* [*j*]) *− gap*_*G*_(*M* [*β*_*x*_], *M* [*j*]). Therefore, *C*_*x*+1_[*j*] ≥ *f*_*x*+1_(*j*).
▪ Case 2: *rank*(*M* [*β*_*x*_].*v*) *> rank*(*M* [*j*].*v*). In this case, *C*[*j*] may not meet the desired threshold after *M* [*j*].*v* is processed because *M* [*β*_*x*_].*v* is processed later than *M* [*j*].*v*. However, *M* [*j*].*v* must be reachable from *M* [*β*_*x*_].*v* using walks through *{last*2*reach*(*M* [*j*].*v, i*) : *i ∈ paths*(*M* [*β*_*x*_].*v*)*}*. Therefore, *C*[*j*] gets updated again due to tuples created in Line 12. This will ensure that *C*_*x*+1_[*j*] ≥ *f*_*x*+1_(*j*).
▪ Case 3: *rank*(*M* [*β*_*x*_].*v*) = *rank*(*M* [*j*].*v*) and *M* [*β*_*x*_].*y < M* [*j*].*x. rank*(*M* [*β*_*x*_].*v*) = *rank*(*M* [*j*].*v*) implies *M* [*β*_*x*_].*v* = *M* [*j*].*v*. The ordering of tuples based on *pos* in Line 21 ensures that the value of key *M* [*β*_*x*_].*d* gets updated in search trees, and *C*[*j*] gets updated afterward.
▪ Case 4: *rank*(*M* [*β*_*x*_].*v*) = *rank*(*M* [*j*].*v*) and *M* [*β*_*x*_].*y* ≥ *M* [*j*].*x*. The tuples created in Line 15 ensure that *C*[*j*] is updated again after finishing the processing of vertex *M* [*j*].*v*. In this case, the gap between anchors *M* [*β*_*x*_] and *M* [*j*] is computed by considering the distance of vertex *M* [*j*].*v* to itself, i.e., 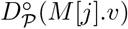.

Accordingly, the time complexity of Algorithm 3 is *O*(Γ_*c*_*N* |𝒫|log *N* + *N* |𝒫|log *N* |𝒫|). In our experiments, we will highlight that parameters Γ_*c*_ and |𝒫|are small in practice. The space complexity of the algorithm is *O*(*N* |𝒫|+ |*V* ||𝒫|) due to construction of array *Z*, the sorting operation on array *Z*, |𝒫|search trees and the precomputed data structures. Next, we show that *O*(*N*) upper bound on the number of iterations is tight.

#### ▸ Lemma 10.

*The count of iterations required by Algorithm 3 is* Ω(*N*) *in the worst-case*.

Proof. An example where Algorithm 3 requires Ω(*N*) iterations is shown in Figure 4. The graph has two vertices forming a cycle. Assume that *weight* of all *N* input anchors is equal and sufficiently high to outweigh the gap cost between any pair of anchors. As *M* [1] *≺ M* [2] *≺ M* [3] … *≺ M* [*N*], the sequence of anchors in the optimal chain is (*M* [1], *M* [2], *M* [3], …, *M* [*N −* 1], *M* [*N*]). After the first iteration of the algorithm, the size of the highest scoring chain computed until then will be 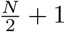. The size will grow slowly by one in each subsequent iteration. A step-by-step dry run of the algorithm is left for the journal version of this paper. ◂

**Figure 4.**
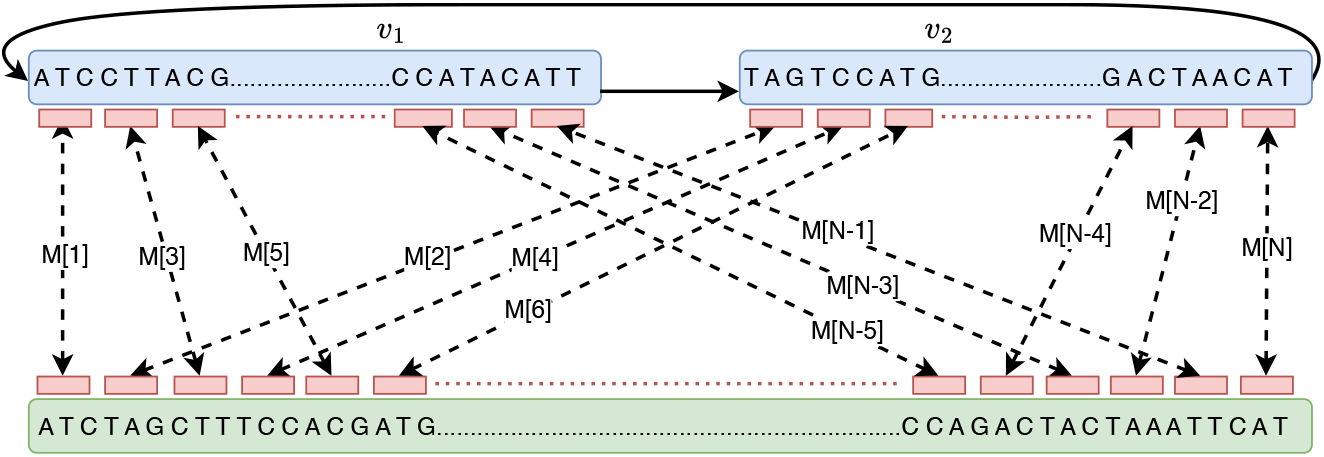
A worst-case example for Algorithm 3 where it requires Ω(*N*) iterations to converge.

## 4 Implementation

We have implemented the proposed algorithm in C++ (https://github.com/at-cg/PanAligner). We call our software as PanAligner. PanAligner is developed as an end-to-end long-read aligner for cyclic pangenome graphs. We borrow open-source code from Minichain [5], Minigraph [24], and GraphChainer [30] for other necessary components besides co-linear chaining. While using PanAligner, a user needs to provide a graph (`GFA` format) and a set of reads or contigs (`fasta` or `fastq` format) as input. We use the standard data structure to store the pangenome graph while accounting for double stranded nature of DNA sequences.

For each vertex *v ∈ V*, we also add another vertex 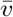 whose string label is the reverse complement of string *σ*(*v*). For each edge *u → v ∈ E*, we add the complementary edge 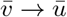. This enables read alignment irrespective of which strand the read was sequenced from. For the benchmark, we built pangenome graphs by using Minigraph v0.20 [24]. Minigraph augments large insertion, deletion, and inversion variants into the graph while incrementally aligning each input assembly. Inversion variants can introduce cycles in the graph because Minigraph augments them by linking the vertices from opposite strands. The graph contains multiple weakly connected components because the components corresponding to different chromosomes are never linked during graph construction. Similar to [5, 30], we consider each weak component independently during both the preprocessing and co-linear chaining stages to enable efficient multithreading and memory optimization.

We defined our problem formulation to produce an optimal chain, but we actually compute multiple best chains, similar to [5, 23, 24]. This is because there can be multiple high-scoring alignments of a read on the graph. PanAligner also outputs a mapping quality score between 0 to 60 to indicate the confidence score for each alignment [25]. We used seeding and extension code from Minigraph [24]. Seeding is done by identifying minimizer matches [45] between vertex labels of the graph and the read. The extension code produces the final base-to-base alignment by joining the chained anchors [52]. We used code from GraphChainer [30] to compute the minimum path cover of a DAG and range queries.

## 5 Experiments

### Benchmark Datasets

We constructed four cyclic pangenome graphs by using subsets of publicly available 95 haplotype-resolved human genome assemblies [26, 37]. These graphs were generated using Minigraph v0.20 [24]. We used CHM13 human genome assembly [37] as the starting sequence during graph construction in all four graphs. We refer to these graphs as 10H, 40H, 80H, and 95H, where the prefix integer represents the count of haplotypes in each graph. The properties of these graphs are provided in Table 1.

**Table 1.** Properties of four cyclic pangenome graphs that were used for evaluation.

| Graph | \|V\| | \|E\| | No. of weak components | No. of structural variants | N50 length of vertex labels (kb) |
| --- | --- | --- | --- | --- | --- |
| 10H | 283,296 | 406,292 | 30 | 61,523 | 225 |
| 40H | 679,846 | 978,122 | 28 | 149,163 | 127 |
| 80H | 1,106,286 | 1,594,980 | 26 | 244,372 | 85 |
| 95H | 1,224,853 | 1,765,222 | 26 | 270,888 | 79 |

### Evaluation Methodology

We simulated long reads using PBSIM2 v2.0.1 [39] from CHM13 assembly with N50 length 10 kb, 0.5*×* sequencing coverage and 5% error-rate to approximately mimic the properties of long-reads. We labeled the IDs of the simulated reads with their true interval coordinates in the CHM13 assembly for correctness evaluation. To confirm the correctness of a read alignment, we used similar criteria from prior studies [5, 23, 24]. We require that the reported walk corresponding to a correct alignment should only use the vertices corresponding to the CHM13 assembly in the graph, and it should overlap with the true walk. We used `paftools` [23] to automate this evaluation. By default, it requires the overlapping portion to be at least 10% of the union of the true and the reported walk length. We executed all experiments on a computer with AMD EPYC 7763 64-core processor and 512 GB RAM. We ran each aligner using 32 threads to leverage the multi-threading capabilities of the tested aligners.

All aligners process reads in parallel. We used the `/usr/bin/time -v` command to measure wall clock time and peak memory usage.

### Size of Path Cover and Count of Iterations

Finding a suitable path cover 𝒫 of the input graph such that |𝒫|*≪* |*V* |is a crucial step in our proposed framework because the scalability of our algorithms depends on this parameter. We discussed a heuristic to compute path cover in Section 3.1 because determining minimum path cover in general graphs is *NP* -hard. Table 2 shows the sizes of path covers computed by our heuristic in all four graphs. Recall that our algorithms process the weakly connected components of a graph independently. In each graph, we indicate the size of the path cover as a range because path covers vary per component. The results show that our heuristic is effective in finding a small path cover (in the number of paths).

**Table 2.** All four graphs have multiple weakly connected components. Therefore, the size of the identified path cover of each graph is presented as a range. The other columns show the statistics for the number of anchors and the number of iterations used by our iterative algorithms (Algorithms 1, 2, 3). The statistics were gathered while aligning simulated long reads to cyclic pangenome graphs.

| Graph | Size of path cover (min-max) | Number of anchors |  | Number of iterations |  |  |  |  |  |
| --- | --- | --- | --- | --- | --- | --- | --- | --- | --- |
|  |  |  |  | Array *last2reach* |  | Array *D* |  | Chaining |  |
|  |  | Mean | Max | Mean | Max | Mean | Max | Mean | Max |
| 10H | 1-20 | 10,900 | 309,591 | 2.0 | 4 | 2.0 | 5 | 2.3 | 77 |
| 40H | 1-36 | 10,850 | 309,603 | 2.0 | 4 | 2.0 | 5 | 2.4 | 72 |
| 80H | 1-49 | 10,804 | 309,364 | 3.0 | 4 | 3.0 | 5 | 2.4 | 61 |
| 95H | 1-59 | 10,798 | 309,367 | 3.0 | 4 | 3.0 | 5 | 2.4 | 64 |

The number of anchors *N* that were provided as input to the co-linear chaining algorithm varies per read. We report the mean and maximum value in Table 2. Observe that *N* does not change much with increasing haplotype count. Next, we evaluate the count of iterations Γ_*l*_, Γ_*d*_ used by our graph preprocessing algorithms (Algorithms 1-2) and also report them as a range for each graph. These algorithms compute *last*2*reach* and *D* arrays. Observe that the iteration count is significantly smaller in practice than the proven upper limit of |*V* |(Lemma 7). This is because the worst-case situation is not observed in practice. Accordingly, there is minimal time overhead during the preprocessing phase.

The count of iterations Γ_*c*_ required by our chaining algorithm (Algorithm 3) varies per component as well as per read. We collect the iteration count statistics as follows. For a single read, we define the iteration count as the maximum number of iterations used over all components. Based on this definition, we report the average and the maximum count over all reads in Table 2. Observe that the average count is *<* 2.5 using all four graphs. The maximum count is *<* 100. These numbers are again significantly better compared to the upper bound from Lemma 9.

### Alignment of Simulated Reads to Cyclic Graphs

We assessed the performance of PanAligner against two sequence-to-graph aligners, Minigraph v2.20 [24] and GraphAligner v1.0.17b [42], that can handle cycles. Unlike PanAligner, Minigraph and GraphAligner use heuristics to join anchors. Minichain [5] and GraphChainer [30] were excluded from this comparison because they do not support cyclic graphs.

We highlight the accuracy, runtime, and memory usage of different aligners using graphs 10H and 95H in Tables 3 and 4, respectively. Observe that PanAligner outperformed Minigraph and GraphAligner in terms of accuracy, i.e., the fraction of correctly aligned reads. This advantage is even more apparent if low-confidence alignments with mapping quality *<* 10 are ignored. We show the comparison plots in Figure 5. Both PanAligner and Minigraph left a small fraction of reads unaligned. This may be because (i) both methods drop high-frequency minimizer matches, and (ii) they do not consider low-scoring chains for the extension stage. In contrast, GraphAligner achieved higher recall by aligning all reads, but this came at the expense of lower accuracy.

**Table 3.** A comparison of the performance of long-read aligners using the 10H graph. MQ stands for mapping quality.

|  | PanAligner | Minigraph | GraphAligner |
| --- | --- | --- | --- |
| Indexing time (sec) | 96 | **57** | 238 |
| Alignment time (sec) | 2924 | **46** | 4928 |
| Memory usage (GB) | **23.14** | 23.19 | 24.68 |
| Unaligned reads | 1.18% | 2.17% | **0%** |
| Incorrectly Aligned reads | **0.79%** | 1.19% | 1.47% |
| Unaligned reads (MQ$\geq 10$) | 3.51% | 5.85% | **0.78%** |
| Incorrectly Aligned reads (MQ$\geq 10$) | **0.20%** | 0.32% | 0.91% |

**Table 4.**
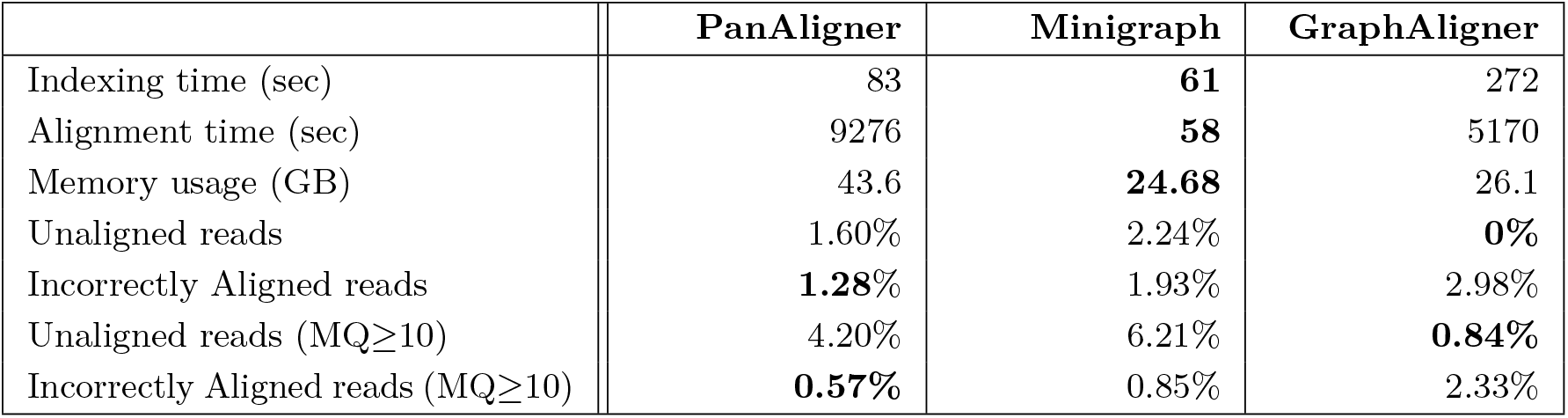
A comparison of the performance of long-read aligners using the 95H graph. MQ stands for mapping quality.

**Figure 5.**
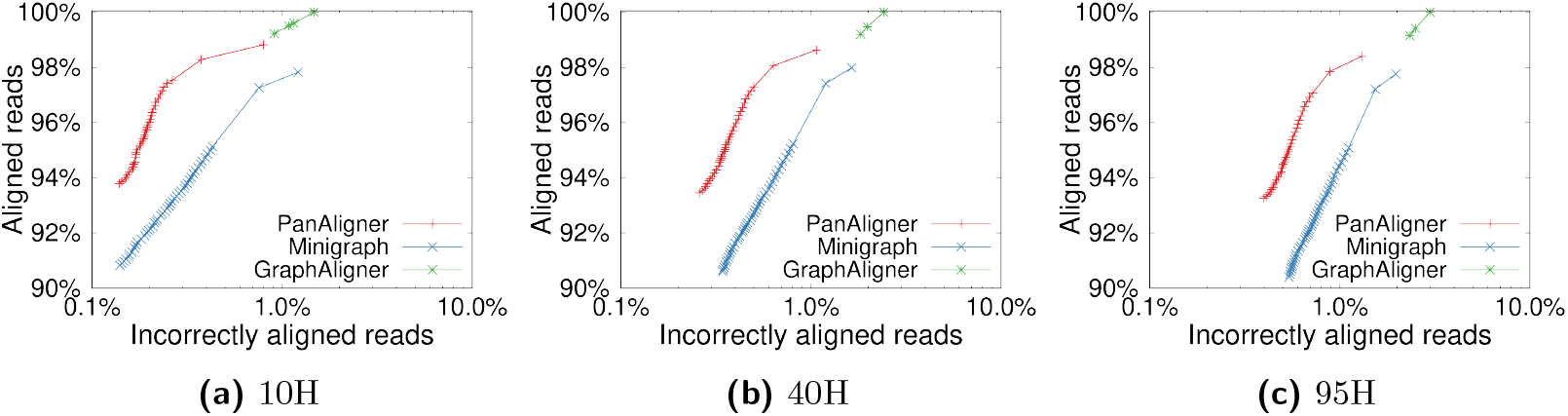
The plots show the fraction of aligned reads and the accuracy obtained by using all the aligners on graphs 10H, 40H, and 95H. These plots were generated by varying mapping quality cutoffs from 0 to 60. X-axis in these plots uses a logarithmic scale to indicate the percentage of incorrectly aligned reads.

Table 2 shows that the size of the path cover computed by our heuristic increases by roughly a factor of three from 10H to 95H. We can see how this parameter proportionally affects PanAligner’s runtime in Tables 3 and 4. PanAligner’s runtime is significantly higher than Minigraph for both 10H and 95H graphs because it uses an iterative algorithm. Runtimes of PanAligner and GraphAligner are in the same order of magnitude. PanAligner’s computational needs are within practical limits, thus making it an effective method for accurately aligning long reads or contigs to cyclic pangenome graphs. We observe a consistent drop in alignment accuracy of all three aligners with increasing haplotype count (Figure 5). This is likely because the number of combinatorial paths to which a read can align grows exponentially with respect to the haplotype count.

### Alignment of Simulated Reads to Acyclic Graphs

We also tested PanAligner for acyclic pangenome graphs. We followed the same procedure as [5] to generate a DAG from 95 haplotype-phased assemblies and refer to this graph as 95H-DAG. This graph was generated by disabling inversion variants during graph construction in Minigraph [24]. 95H-DAG has 1.2M vertices and 1.8M edges. We also include Minichain v1.0 [5] and GraphChainer v1.0.2 [30] in this comparison. GraphChainer uses a co-linear chaining algorithm for DAGs without penalizing gaps. For DAG inputs, the problem formulation in PanAligner becomes equivalent to the one used in Minichain [5]. A single iteration of our algorithms suffices for DAGs. Therefore, we simply check if the input graph is a DAG at the preprocessing stage, and run a single iteration of Algorithms 1-3. PanAligner achieves similar performance as Minichain in terms of speed and accuracy for DAGs (Table 5). It compares favorably to other methods in terms of accuracy.

**Table 5.** A comparison of the performance of long-read aligners using the 95H-DAG graph. MQ stands for mapping quality.

|  | Pan-Aligner | Minichain | Mini-graph | Graph-Aligner | Graph-Chainer |
| --- | --- | --- | --- | --- | --- |
| Indexing time(sec) | 78 | 77 | **62** | 276 | 575 |
| Alignment time(sec) | 2406 | 2515 | **50** | 5136 | 23710 |
| Memory usage (GB) | 30.04 | 25.61 | **24.79** | 26.12 | 185.83 |
| Unaligned reads | 1.62% | 1.62% | 2.23% | **0%** | **0%** |
| Incorrectly Aligned reads | **1.28%** | 1.29% | 1.92% | 3.06% | 4.93% |
| Unaligned reads (MQ≥10) | 4.75% | 4.75% | 6.26% | 0.85% | **0%** |
| Incorrectly Aligned reads (MQ≥10) | **0.53%** | 0.54% | 0.84% | 2.41% | 4.93% |

## 6 Discussion

Co-linear chaining is a fundamental technique for scalable sequence alignment. Several classes of structural variants, such as duplications, tandem repeat polymorphism, and inversions, are best represented as cycles in pangenome graphs [41, 26]. Existing alignment software designed for cyclic graphs are based on heuristics to join anchors [24, 42]. We proposed the first practical problem formulation and an efficient algorithm for co-linear chaining on pangenome graphs with cycles. We gave a rigorous analysis of the algorithm’s time complexity. The proposed algorithm serves as a useful generalization of the previously known ideas for DAGs [5, 29, 30, 33]. We implemented the proposed algorithm as an open-source software PanAligner. We demonstrated that PanAligner scales to large pangenome graphs built by using haplotype-phased human genome assemblies. It offers superior alignment accuracy compared to existing methods.

In our formulation, we did not allow anchors to span two or more vertices in a graph for simplicity of notations, but the proposed ideas can be generalized. PanAligner software borrows seeding logic from Minigraph [24], which also restricts anchors within a single vertex. This simplification is appropriate if the graph only includes structural variants (*>* 50 bp). The current version of PanAligner software may not be suitable for graphs which include substitution and indel variants.

Future work will be directed in the following directions. First, we will test the performance of PanAligner on pangenome graphs that are constructed by using alternative methods, e.g., [16, 19, 26]. Second, we will explore formulations for haplotype-constrained co-linear chaining to control the exponential growth of combinatorial search space with the increasing number of haplotypes [34, 48]. Third, we will generalize the proposed techniques for aligning reads to long-read genome assembly graphs which also contain cycles. It will be interesting to understand whether the small width assumption is appropriate for assembly graphs.

## Acknowledgements

The authors thank Manuel Cáceres, Shravan Mehra and Sunil Chandran for providing useful feedback.

